# Quantifying collective interactions in biomolecular phase separation

**DOI:** 10.1101/2023.05.31.543137

**Authors:** Hannes Ausserwöger, Daoyuan Qian, Georg Krainer, Ella de Csilléry, Timothy J. Welsh, Tomas Sneideris, Titus M. Franzmann, Seema Qamar, Nadia A. Erkamp, Jonathon Nixon-Abell, Mrityunjoy Kar, Peter St George-Hyslop, Anthony A. Hyman, Simon Alberti, Rohit V. Pappu, Tuomas P. J. Knowles

**Affiliations:** Centre for Misfolding Diseases, Yusuf Hamied Department of Chemistry, University of Cambridge, Lensfield Road, Cambridge CB2 1EW, United Kingdom; Biotechnology Center (BIOTEC), Center for Molecular and Cellular Bioengineering (CMCB), Technische Universität Dresden, Tatzberg 47/49, Dresden, Germany; Cambridge Institute for Medical Research, Department of Clinical Neurosciences, Clinical School, University of Cambridge, Cambridge, CB2 0XY, United Kingdom; Max Planck Institute of Cell Biology and Genetics (MPI-CBG), 01307 Dresden, Germany; Department of Medicine (Division of Neurology), Temerty Faculty of Medicine, University Health Network, University of Toronto, Toronto, Ontario M5T 0S8, Canada; Department of Neurology, Columbia University, 710 West 168th Street, New York, New York 10032, USA; Department of Biomedical Engineering and Center for Biomolecular Condensates, Washington University in St. Louis, St. Louis, MO, USA; Cavendish Laboratory, Department of Physics, University of Cambridge, JJ Thomson Road, Cambridge CB3 0HE, United Kingdom

**Author notes:** **Correspondence** Correspondence and requests for materials should be addressed to Tuomas P. J. Knowles. Contributed equally (Hannes Ausserwöger, Daoyuan Qian, Georg Krainer).

## Abstract

Biomolecular phase separation plays a pivotal role in governing critical biological functions and arises from the collective interactions of large numbers of molecules. Characterising the underlying collective interactions of phase separation, however, has proven to be challenging with currently available tools. Here, we propose a general and easily accessible strategy to quantify collective interactions in biomolecular phase separation with respect to composition and energetics. By measuring the dilute phase concentration of one species only, we determine tie line gradients and free energy dominance as dedicated descriptors of collective interactions. We apply this strategy to dissect the role of salts and small molecules on phase separation of the protein fused in sarcoma (FUS). We discover that monovalent salts can display both exclusion from or preferential partitioning into condensates to either counteract charge screening or enhance non-ionic interactions. Moreover, we show that the common hydrophobic interaction disruptor 1,6-hexanediol inhibits FUS phase separation by acting as a solvation agent capable of expanding the protein polypeptide chain. Taken together, our work presents a widely applicable strategy that enables quantification of collective interactions and provides unique insights into the underlying mechanisms of condensate formation and modulation.

## Introduction

The assembly of proteins, driven by biomolecular interactions, is a ubiquitous phenomenon in living cells that enables essential functions spanning from protein synthesis^1,2^ to cellular signalling^3,4^. Recently, the phase separation of proteins into biomolecular condensates has emerged as a new theme for protein assembly^5–7^. Through phase separation, proteins demix from a homogenous phase into a protein-rich dense phase and a protein-poor dilute phase^8,9^. This process of condensate formation has been shown to be critical in diverse biological processes including gene expression and cellular organisation^10–13^, and its misregulation has been associated with the emergence of aberrant cellular functions and diverse pathologies^14^.

Biomolecular phase separation is driven by collective interactions, stemming from the collaborative association of large numbers of molecules. These collective interactions give rise to complex emergent behaviours, which demand dedicated descriptors for their quantification. This includes characterizing the partitioning of species between dense and dilute phases and understanding the individual components’ contributions to the decrease in free energy. Common protein interaction assays fail to inform on the collective properties of the interactions driving phase separation. This is because traditional approaches are centred around determining binding energetics between individual molecules rather than large collectives. Even assays that map out the phase diagram and thereby characterise the critical concentrations of the two-phase coexistence region^15–17^, do not provide information on condensate composition and fail to quantify the energetic driving forces.

Recent theoretical advancements have introduced novel descriptors that enable quantification of collective interactions^18,19^. These descriptors are derived from the analysis of tie line gradients^18^ and the quantification of component dominance^19^, and provide means to physically characterise the mechanisms of condensate formation and modulation. Tie lines describe the demixing process between biomolecular dense and dilute phases, providing insights into condensate composition and the underlying interactions. Additionally, component dominance analysis offers the opportunity to discern the individual components’ contributions to the decrease in free energy, thus shedding light on the underlying driving forces and energetics in a locally defined manner. Hence, translating these theoretical advances into experiments would enable to addres the current limitations in characterising the collective interactions underlying protein phase separation.

Here, we present a general and broadly applicable strategy for the experimental characterisation of the collective interactions underlying biomolecular phase separation. This strategy is based on determining tie line gradients and the analysis of component dominance, which enables quantification of partitioning, stoichiometry, and the relative free energy contributions of individual components in a phase separating system. Experimentally, this is achieved by measuring the dilute phase concentration of a single component only. We apply this strategy to gain insights into the collective interactions of the phase separating protein fused in sarcoma (FUS) under the effect of salts at low and high ionic strength conditions and to study the modulation of FUS condensates by the small-molecule hydrophobic disruptor 1,6-hexanediol.

### Quantifying collective interactions

Tie lines describe the compositions formed after demixing of a given set of total concentrations (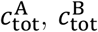, etc. for components A, B, etc.) within the coexistence region of the phase diagram. More specifically, tie lines connect the equilibrium dilute and dense phase concentrations of constituent species (Fig. 1a), including the partitioning of all species in the system, such as salts and buffer excipients. These higher dimensional tie lines can be approximated by the so-called reduced tie lines, based on the dimensionality reduction of the component A and B measurement plane of interest. Crucially, the reduced tie line gradient *K* preserves essential information on partitioning of (co-)solutes of interest as it approximates the ratio of the higher dimensional tie line gradient entries in A and B (see Supplementary Information and Supplementary Fig. S1 for details on tie line reduction)^19^. Specifically, for a negative reduced tie line gradient (***K*** < 0), component A is enriched in the dense phase (i.e., included in condensates), while component B is decreased (i.e., excluded from condensates). By contrast, a positive reduced tie line gradient (***K*** > 0) means that both species are present at higher concentrations in the dense phase than in the dilute phase, indicating preferential partitioning into condensates.

**Figure 1.**
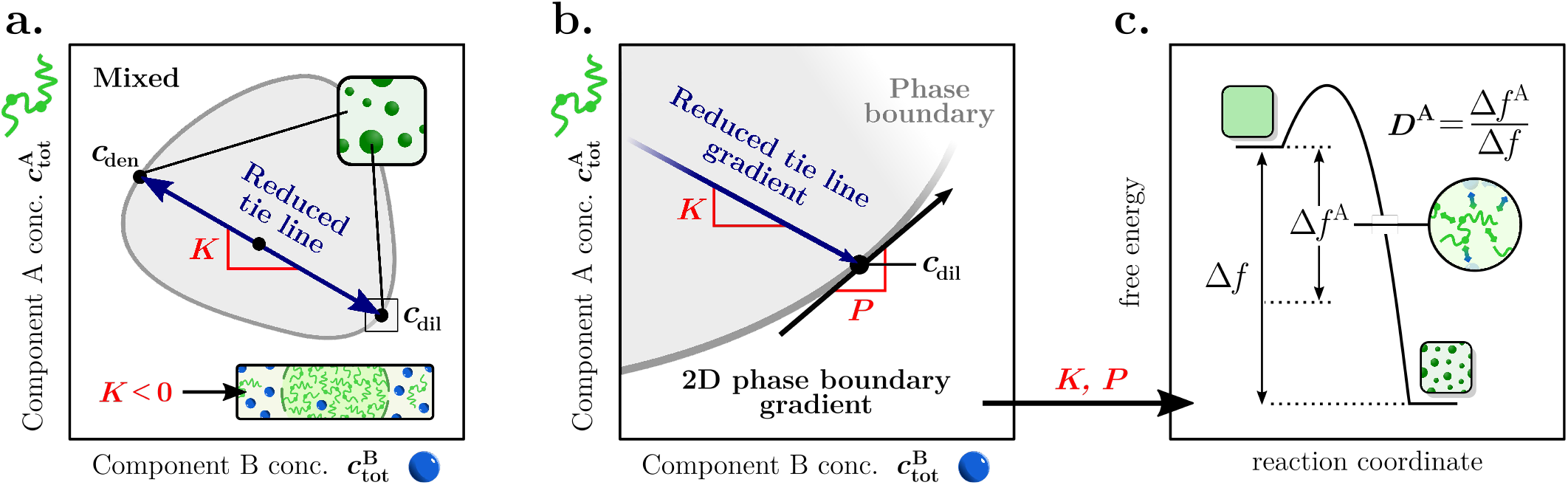
Descriptors of collective interactions in biomolecular phase separation. **(a)** Schematic of a full phase diagram for a 2D system between component A and B, where total component A and B concentrations 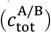 within the coexistence region are shaded in grey. Tie lines describe the demixing within the coexistence region into dilute (*c*_dil_) and dense phase concentrations (*c*_den_). The reduced tie line gradient *K*, which is constrained to the 2D measurement plane of interest informs on partitioning. **(b)** Locally, the dependence of the saturation concentration on both component A and B is described by the phase boundary gradient *P*. **(c)** Together the reduced tie line gradient *K* and the local phase boundary gradient *P* inform of the relative free energy decrease for phase separation of component A (Δ*f* ^A^/Δ*f*) as given by the dominance metric of component A (*D*^A^).

In addition to tie lines, the shape of the phase boundary plays an essential role in understanding collective interactions. The phase boundary describes the local dependence of the saturation concentration (Fig. 1b) on the individual components of interest. This relationship can be quantified by the local phase boundary gradient *P*. Exemplary, in the extreme cases where *P* approaches 0 or ∞, the onset of phase separation becomes solely dependent on component B or A, respectively. Together, the local phase boundary gradient *P* and the reduced tie line gradient *K* together inform on the energetics of the phase separation process according to (Fig. 1c)^19^:

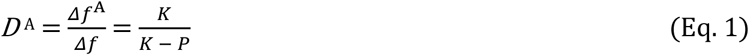

where *D*^A^ denotes the dominance of component A and *Δf*^A^ the free energy decrease of component A with regards to the overall free energy decrease Δ*f*^19^. Hence, *D*^A^ gives the relative contribution of component A to the overall free energy decrease, which provides a measure of the systems energetic dependence on said component and allows for the characterisation of the underlying energetic driving forces of phase separation in a locally defined manner.

Crucially, reduced tie line gradients *K*, phase boundary gradients *P* and dominance *D*^A^ are easily determinable from measuring dilute phase concentrations of one component only, as shown in Figure 2. In our approach, dilute phase concentrations are measured experimentally by injecting samples into a microfluidic device and reading out the fluorescent signal of one of the species of interest (i.e., protein = component A) by confocal detection (see Fig. 2a; Supplementary Fig. S2 for confocal measurement set-up). The dilute phase concentrations are extracted from the baseline signal in the fluorescence time traces. The presence of the dense, condensate phase is detected as short bursts of high photon intensities stemming from high protein concentrations within condensates (Fig. 2b and c). Although confocal based measurements in conjunction with microfluidics provide high precision and throughput, the approach can be similarly applied with any other means of dilute phase concentration readouts such as centrifugation combined with standard epifluorescence or absorbance-based measurements.

**Figure 2.**
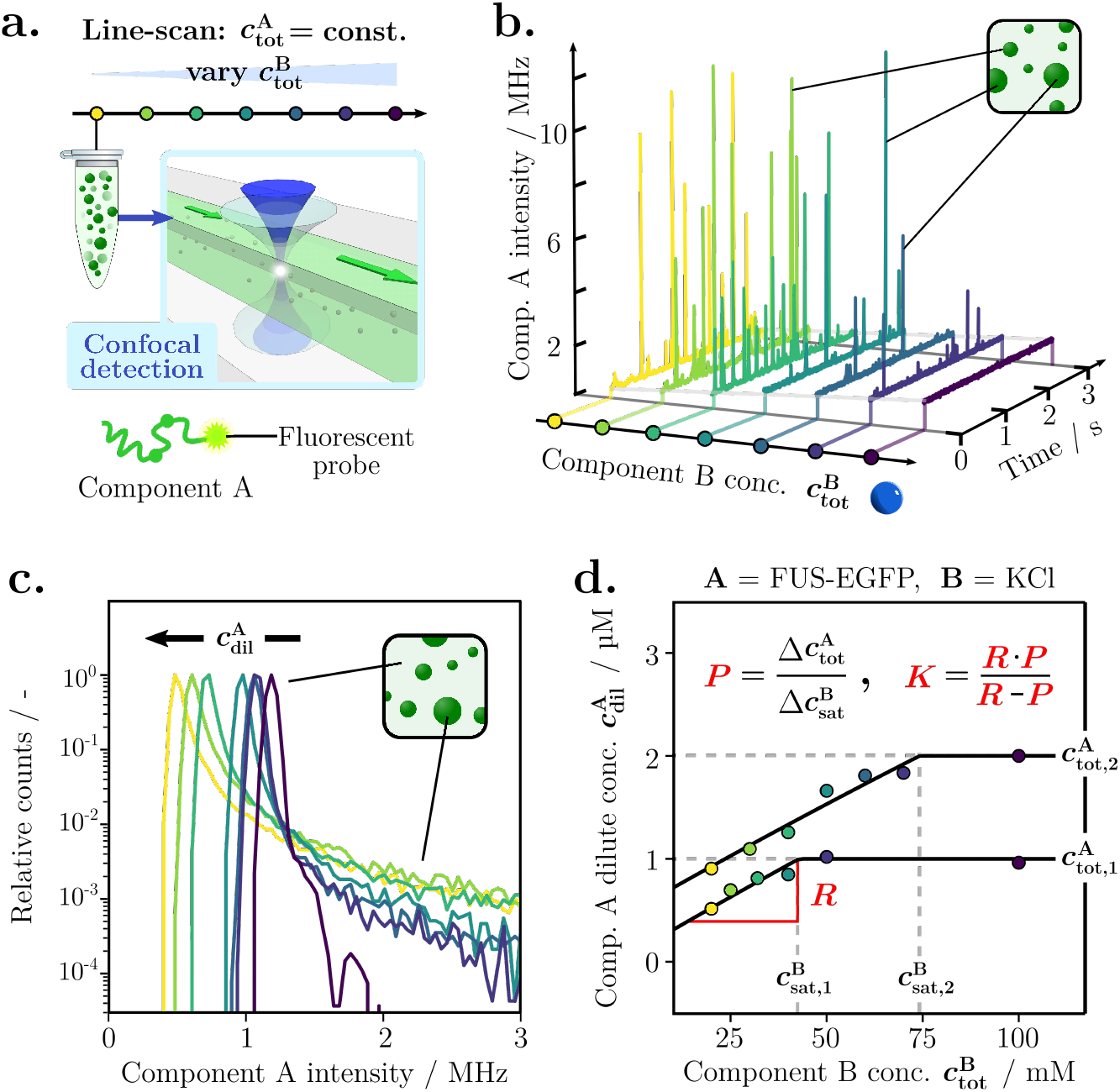
Quantification of collective interactions using one component dilute phase concentrations measurements. **(a)** Line scans are performed by injecting samples with constant concentrations of component A 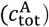 but varying concentrations of component B 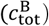 into a microfluidic flow cell. Readout of fluorescence is done by confocal detection. **(b)** Intensity time traces of component A against varying 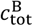 obtained from confocal detection measurements. **(c)** Intensity histograms obtained from time traces in (b), where the maximum is representative of the dilute phase concentration of component A. **(d)** Component A (FUS-EGFP), dilute phase concentration 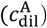 changes with varying component B (KCl) concentrations along two separate line scans. The raw data for the line scan at 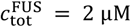 is shown in (b, c) and the corresponding data for the line scan at 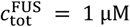 is given in Supplementary Fig. S3. Evaluation of compositional and thermodynamic parameters follows from the dilute phase response gradient *R* and the saturation concentrations.

To extract the aforementioned parameters, the dilute phase concentration measurements are performed in the form of so-called line scans. The change in the dilute phase concentration of component A is quantified along a series of experiments with a constant total concentration of component A 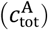 while adjusting the total concentration of component B 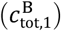. Two sets of such line scans at two different constant total concentrations of component A (i.e., 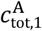 and 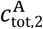 ) are required to extract the relevant parameters. Figure 2d illustrates typical line scan measurements. At higher concentrations of component B, the line scan displays a flat plateau region because no phase separation occurs in this range. Consequently, the dilute phase concentration is equal to the total concentration. However, at total concentrations of component B below the saturation concentration of component B 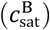, phase separation takes place, leading to a decrease in the dilute phase concentration due to the formation of condensates.

From the obtained dilute phase line scans, the parameters *P, K* and *D*^A^ are extracted. The phase boundary gradient *P* is determined from the change in 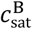 with changing 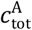 :

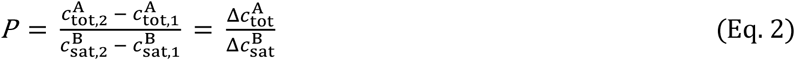

In order to calculate the reduced tie line gradient *K*, the dilute phase response gradient *R* is determined from the initial slope of the decrease in dilute phase concentration (Fig 2d, see Materials and Methods for details). *R* quantifies the change in dilute phase concentration of component A with increasing concentrations of component B. From *P* and *R* the reduced tie line gradient *K* is calculated according to^19^:

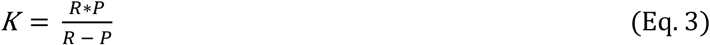

Finally, from *K* and *P*, the dominance *D* of component A is determined according to Eq. 1, yielding a full set of parameters that enable the description of collective interactions of biomolecular condensates. In the following, we apply this approach for compositional and energetic quantification of collective interactions underpinning phase separation and modulation of the protein FUS.

### Ions are excluded from FUS condensates at low ionic strengths to counteract charge screening

The intrinsically disordered protein FUS is involved in physiologically and pathologically relevant intracellular condensation events^13,20–22^, the emergence of amyotrophic lateral sclerosis (ALS)^13,23,24^, and various forms of cancer^25,26^. One of its hallmarks is its tendency to undergo phase separation at low ionic strengths. However, fundamental features such as ion partitioning or the impact of ions on the energetic driving force remain unaddressed. To address this challenge, we set out to characterise the collective interactions of FUS with salt species.

In a first set of experiments, we examined FUS phase separation at low concentrations of potassium chloride (KCl) (Fig. 3a). Specifically, we measured the dilute phase concentrations of FUS 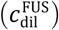 at 1 and 2 μM total protein concentration (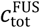, FUS = component A) and KCl concentrations (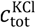, KCl = component B) ranging from 20 to 150 mM (Fig. 3b and Supplementary Fig. S3). At high KCl concentrations (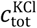 > 100 mM), 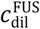 was constant as the protein did not undergo phase separation (see Fig. 3b). Only with decreasing charge screening at lower KCl concentrations (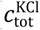 ∼ 40–75 mM, depending on protein concentration), a drop-off in 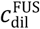 and the appearance of high-intensity bursts were observed, corresponding to the onset of phase separation. This enables determination of the dilute phase response gradient *R* = 0.020 ± 0.003 μM/mM, which indicates that increasing KCl concentrations cause an increase in the FUS dilute phase concentration. The phase boundary gradient of *P* = 30.3 ± 8.1 μM/mM highlights the dependence of phase separation onset and KCl concentration. Specifically, much lower concentrations of KCl are needed to still induce phase separation at lower protein concentrations. This can be further illustrated by determining the FUS/KCl phase boundary in the experimentally probed concentration range using the dilute phase concentration responses and the reduced tie line gradients (Fig. 3c and Supplementary Fig. S4 for details).

**Figure 3.**
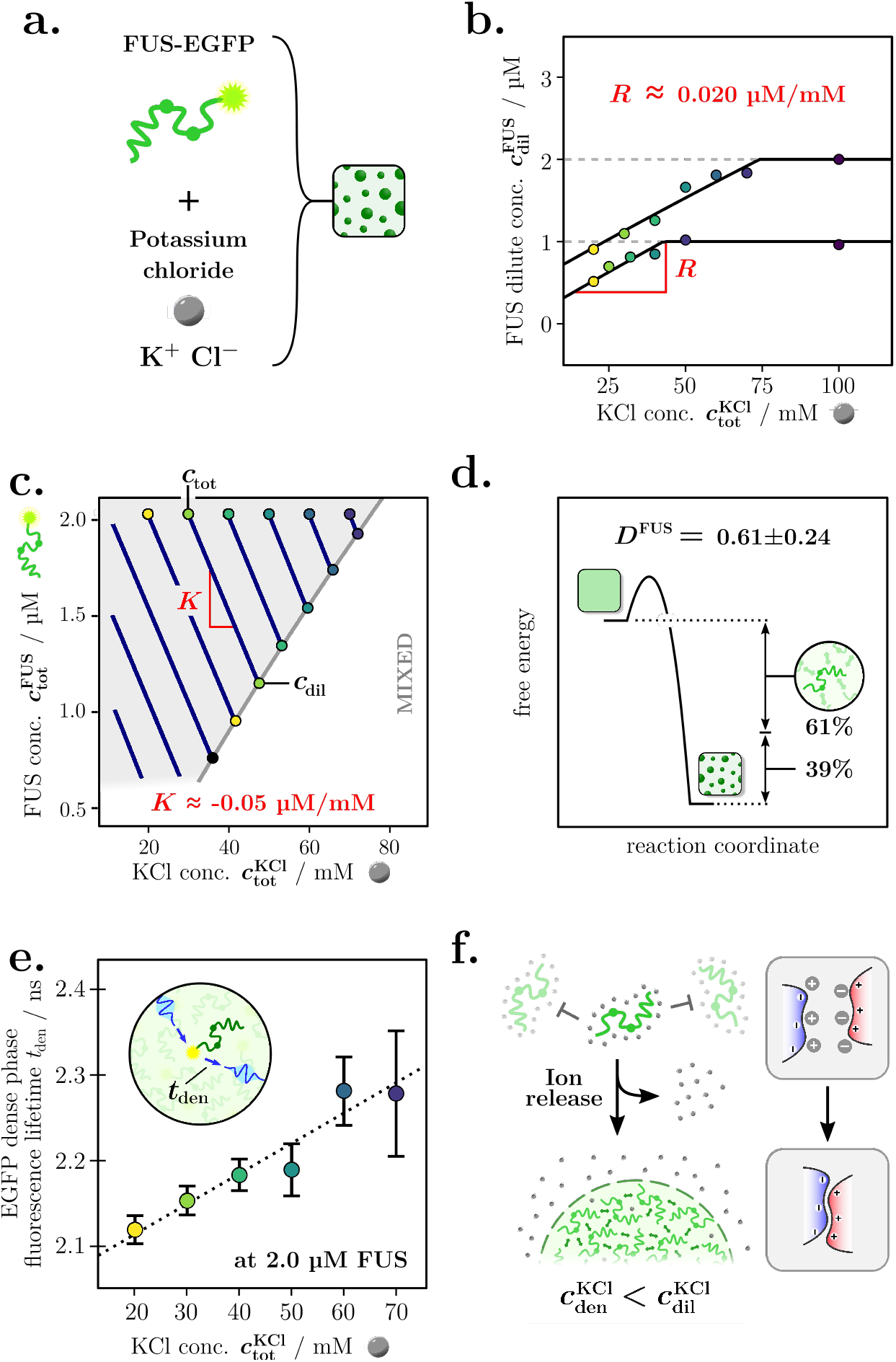
Ion release drives FUS phase separation at low salt conditions. **(a)** Phase separation of FUS-EGFP was studied in the presence of potassium chloride (KCl). **(b)** Dilute phase concentration changes of FUS 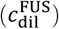 at varying total KCl concentrations 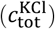. **(c)** Determination of the FUS/KCl phase boundary from dilute phase concentration changes in (**b**) (see Materials and Methods for details). **(d)** Only 61% of the free energy change from phase separation is associated with FUS as per dominance metric analysis (*D*^A^). **(e)** Dense phase EGFP fluorescent lifetimes (*t*_den_) at 2 μM FUS and varying KCl concentrations. **(f)** Schematic summary of ion release mechanism driving FUS phase separation at low KCl concentrations.

The compositional details of the collective FUS–KCl interaction can then be assessed by determining the reduced tie line gradient ***K*** = – 0.05 ± 0.04 μM/mM. Here, considering the dissociation of the salt into K^+^ and Cl^-^, the obtained gradient corresponds to the ratio of protein to a weighted sum of the cationic and anionic species, with the weights relating to their valency^19^. *K* < 0 shows that the ion concentration is lower in the dense phase than in the dilute phase (Fig. 3c). Hence, on average the ions are preferentially excluded from FUS condensates as previously indicated by simulation results for FUS condensates^27^. Preferential exclusion of ions from the condensed phase has also been observed for polyelectrolyte polymers^28,29^.

The exclusion of ions from the condensed phase is particularly interesting as simulations indicate that KCl has a propensity to interact with a large fraction of available polypeptide side chains^30^. The ions form a hydration shell around the dispersed protein, which inhibits intermolecular protein interactions. A release of ions from the hydration shell would therefore allow for FUS to engage in much more effective protein–protein interactions within condensates. Indeed, we determined a FUS dominance of *D*^FUS^ = 0.61 ± 0.24, meaning that only ∼61% of the free energy decrease released by phase separation can be associated with FUS (Fig 3e). This highlights the importance of the ion release mechanism from polypeptide chains to enable protein–protein interactions within condensates to the overall energetic driving force.

The proposed ion release mechanism suggests that the condensate scaffold and the environment itself changes drastically with salt concentration. To investigate this further, we determined the fluorescence lifetime of the EGFP protein tag on FUS as a function of varying KCl concentrations. Distinctly different lifetimes were observed for the dilute phase and the condensate phase (*t*_dil_ ∼ 2.727 vs *t*_den_ < 2.35 ns, changing with KCl concentration, see Supplementary Fig. S5). The much shorter lifetime of the dense phase indicates a higher refractive index due to the locally higher protein concentration within the condensates^31,32^. Crucially, the fluorescence lifetime of the condensed phase was determined to decrease further with decreasing KCl concentration (*t*_den, 70 mM KCl_ = 2.28 ± 0.07 ns, *t*_den, 20 mM KCl_ = 2.12 ± 0.02 ns, both at 2 μM FUS) (see Fig. 3f). Hence, FUS condensates become more densely packed, which is indicative of enhanced protein interactions through the removal of potassium chloride ions. This further supports our observations above that ion release from the FUS poly-peptide chain acts as the key trigger for phase separation (Fig. 3g).

### Ions partition into condensates at high salt concentrations to drive non-ionic interactions

FUS, as well as other proteins, has also been shown to display phase separation at high salt concentrations within a so-called reentrant phase separation regime^33^ (Fig. 4a). To further investigate the mechanism by which ions can trigger phase separation in the high concentration reentrant regime, we performed dilute phase concentration measurements of FUS in the presence of high concentrations of lithium chloride (LiCl) and caesium chloride (CsCl). Both LiCl and CsCl have been previously shown to display large differences in the required saturation concentration^33^, however, the mechanistic basis of this behaviours has remined elusive.

**Figure 4.**
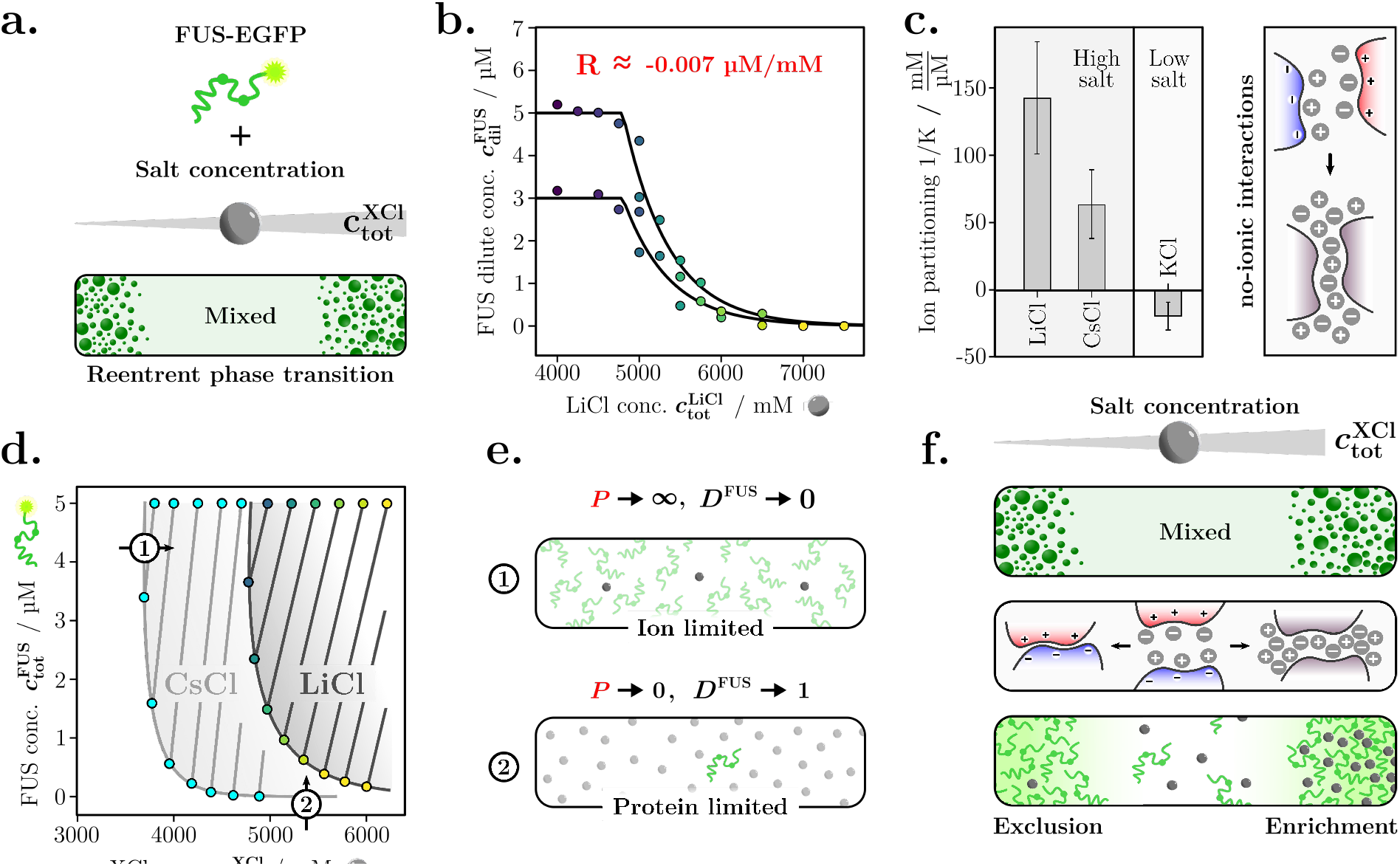
Salt ions are enriched in high salt reentrant FUS condensates to foster non-ionic interactions. **(a)** FUS displays reentrant phase separation at high salt concentrations. **(b)** FUS dilute phase line scans at varying total LiCl concentrations 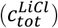. **(c)** Comparison of ion partition (1/*K*) of different salts with respect to FUS condensates. Preferential partitioning at high ionic strengths drives enhancement of non-ionic interactions. **(d)** Phase boundaries show that phase separation occurs only above the respective critical salt and FUS concentrations. **(e)** This highlights dominance extreme cases where the phase separation is driven by only the limiting component. **(e)** Schematic illustration summarizing the mechanism of phase separation in the low and high salt regime.

We first characterised the change in dilute phase concentration of FUS-EGFP under varying concentrations of LiCl at 3 and 5 μM total protein concentration 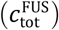 and LiCl concentrations varying between 4 and 7.5 M (Fig. 4b). Indeed, a rapid drop-off in 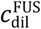, indicated by the onset of reentrant phase separation, was observed at around 5 M LiCl (see Supplementary Fig. S6 for time traces and intensity histograms), consistent with previous observations^33^. From the changes in 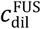 with varying LiCl concentrations, we then determined the ion partitioning of LiCl into FUS condensates from the reduced tie line gradient as 1/*K* = 143 ± 48 mM/μM. Therefore, LiCl at high salt concentrations displays preferential partitioning into condensates. Considering that the drastic increase in ionic strength brings forth strong non-ionic interactions between the protein molecules, the recruitment of ions into the condensed phase will further strengthen this effect. Hence, in the high salt reentrant regime, the salt displays an opposing dense phase partitioning trend compared to the low ionic strength regime (Fig 4c).

Next, we characterised changes in 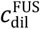 as a function of CsCl concentrations (Supplementary Fig. S7 and S8 for time traces and intensity histograms) to elucidate the impact of changing the cation from Li to Cs. Phase separation was already observed at approx. 3.8 M CsCl compared to a saturation concentration of around 5 M for LiCl, in accordance with previous observations^33^, indicating that CsCl is more potent at triggering reentrant phase separation of FUS. This is also highlighted by the dilute phase concentration changes, where a much steeper drop-off in 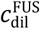 is observed when increasing CsCl (see Fig. 3b and Supplementary Fig. S7). This is further reflected in the ion partitioning for CsCl with FUS, which was determined as 1/*K* = 64 ± 25 mM/μM compared to 1/*K* = 143 ± 48 mM/μM for LiCl. Thereby, CsCl partitions less strongly into the condensed phase, suggesting that less ions are required to allow for sufficient non-ionic interactions occur to trigger phase separation. This is further highlighted by the fact that the tie line gradient appears to be closely linked to the solubility of the salts individually with 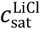 being 13.4 M (1/*K* = 143 mM/μM) and 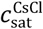 being 5.9 M (1/*K* = 64 mM/μM) (according to supplier specification) as well as following the expected trend of the Hofmeister series^34^.

The reentrant phase transition further highlights extreme cases with regards to the energetics. The salt saturation concentrations at both tested protein concentrations (3 and 5 μM) are very similar for both LiCl and CsCl indicating a sole dependence of phase separation on the salt concentration. Here, the phase boundary gradient approaches *P* → ∞ and thereby *D*^FUS^ → 0, meaning little free energy release is associated with the protein. This is because the system is entirely limited by the ion concentration, highlighting a critical stoichiometry that is necessary to trigger the formation of non-ionic interactions. In addition, 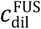 decreased to almost 0 μM protein concentration at high salt concentrations, suggesting that in this ‘salting out’ regime, the protein is almost completely sequestered into the condensed phase. Here, the phase boundary gradient approaches *P* → 0 and with it *D*^FUS^ → 1, suggesting that the system moves from an entirely salt-limited state to a protein-limited state. This can be rationalised by the fact that the salt solubility in the absence of protein is significantly larger.

### 1,6-hexanediol alters the solvation properties of biomolecules to destabilise condensates

Hydrophobic interactions and non-ionic interactions, shown to be critical in driving phase separation in the high salt reentrant regime, are commonly modulated by aliphatic alcohols, such as 1,6-hexanediol (1,6-HD) which causes dissolution of condensates^33,35^. Applying 1,6-HD has become a powerful strategy for studying phase separation to investigate the nature of interactions underlying condensate formation or to assess the liquidity of condensates^36–39^. It is, however, largely unclear how the protein physicochemical properties and collective interactions are altered in the presence of 1,6-HD.

To shed light on the impact of 1,6-HD on collective interactions, we first measured changes in 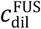 in the absence and presence of 1,6-HD (Fig. 5a, time traces and intensity histograms in Supplementary Fig. S9 and 10). To do so, line-scans were performed at a total FUS concentration of 1 and 2 μM where the total PEG concentration was varied from 0–8 % (w/v) to induce phase separation of FUS. In the absence of 1,6-HD, phase separation of FUS was observed already at a PEG concentration of 2–3 % (w/v) as shown by a decrease in 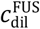 (Fig. 5b). As expected, the addition of 1,6-HD led to an increase in the PEG concentrations necessary to induce phase separation. Accordingly, a decrease of 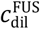 was observed only at around 5–6 % (w/v) PEG. This is also reflected in the shift of the phase boundary, which shows that higher concentrations of PEG are necessary to induce phase separation of FUS in the presence of 1,6-HD (Fig. 5c).

**Figure 5.**
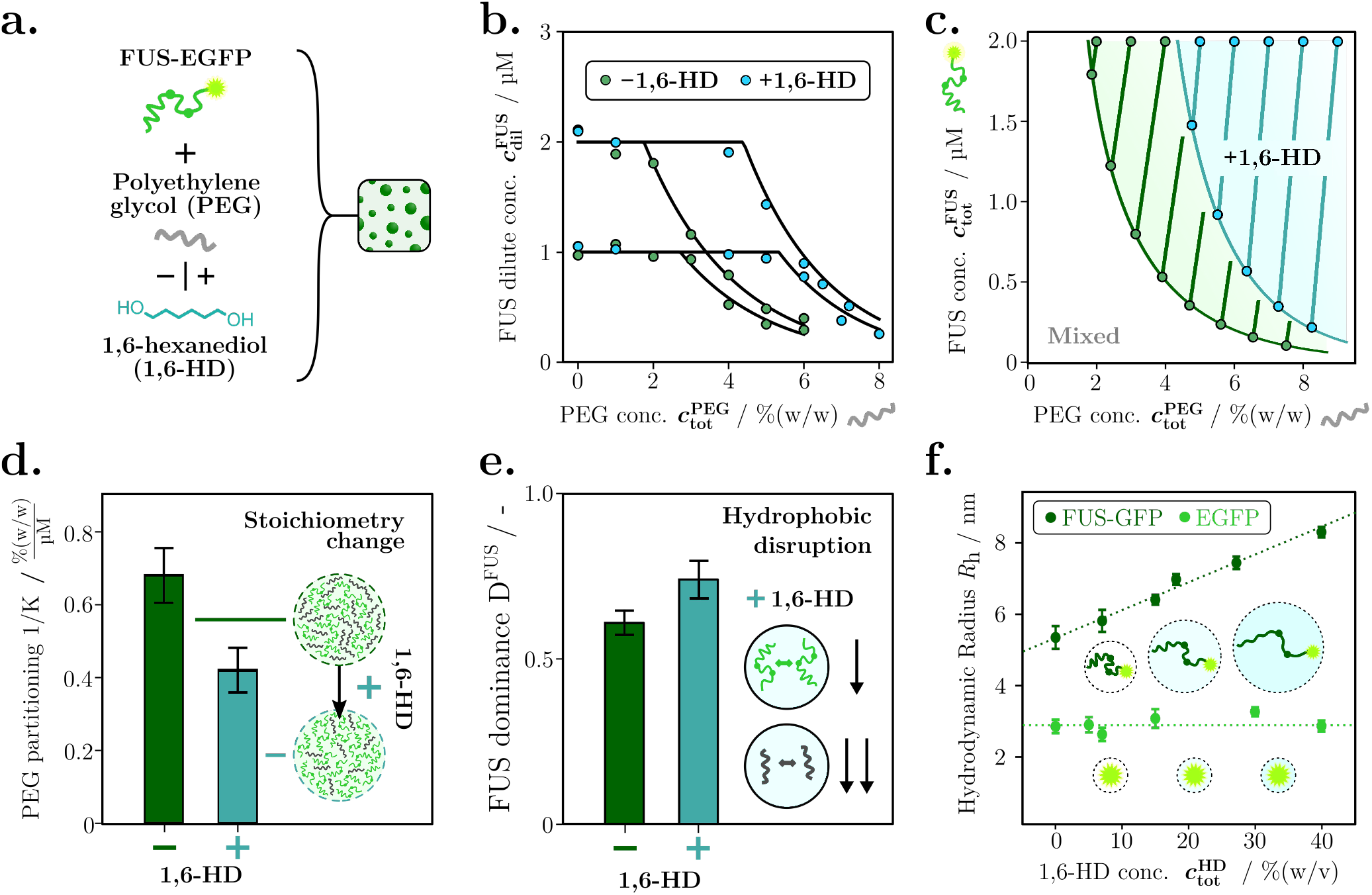
1,6-hexanediol (1,6-HD) decreases the phase separation propensity of FUS by acting as a solvation agent for biomolecules. **(a)** To characterise the mechanism of action of 1,6-hexanediol, phase separation of FUS with PEG was studied in presence (+) and absence (-) of the compound. **(b)** FUS dilute phase concentration changes with varying concentrations of polyethylene glycol (PEG) in the presence (+) and absence (-) of 1,6-HD (7% (w/v)). **(c)** Phase boundaries for FUS against PEG in the absence and presence of 1,6-HD created from changes in dilute phase concentrations and tie line gradients. **(d)** Shift in PEG partitioning (1/K) in the absence and presence of 1,6-HD indicating a change in condensate stoichiometry by decreasing PEG/protein ratio. **(e)** FUS dominance increase in the presence of 1,6-HD indicates that phase separation is suppressed but PEG interactions are affected more strongly. **(f)** Changes in hydrodynamic radius *R*_h_ as a function of 1,6-HD concentration. Shown are data for monomeric FUS-EGFP (dark green) and EGFP protein alone (light green) at 1 μM protein concentration each.

The reduced tie line gradient between FUS and PEG decreases upon addition of 1,6-HD compared to the FUS and PEG system alone (Fig. 5d). Accordingly, PEG partitioning (1/*K*) decreased from 0.69 ± 0.15 % (w/v) / μM without hexanediol to 0.38 ± 0.12 % (w/v) / μM in the presence of 1,6-HD. From the partitioning behaviour, the dense-phase stoichiometry of constituent components can be inferred^18^. The results suggest that the presence of 1,6-HD decreases the number of PEG copolymers per FUS molecule by approximately half within condensates. Such a decrease in copolymer partitioning points towards decrease in interaction strength of PEG. Interestingly, dominance metric analysis shows that the free energy decrease associated with FUS, however, does not change significantly. The dominance of FUS in the absence of 1,6-HD was determined to be *D* ^FUS^ = 0.61 ± 0.08 compared to *D*^FUS^ = 0.74 ± 0.12 in presence of 1,6-HD. This indicates that FUS also experiences a decrease in the self-interaction. This could be explained by 1,6-HD functioning as a solvation agent of the protein by decreasing the solvent polarity. The compositional changes can be traced back to the fact that FUS can also engage in electrostatic interactions, as is well-known from its phase separation propensity at low ionic strengths, while PEG with only ether functional groups will be less capable to do so. FUS, however, is not capable of fully compensating these effects with its electrostatic interactions, which leads to dissolution.

The potentially improved solvation of an intrinsically disordered protein, such as FUS, would be expected to lead to an expansion of its polypeptide chain. Using microfluidic diffusional sizing^40^, we determined the hydrodynamic radius *R*_h_ of monomeric FUS-EGFP at different concentrations of 1,6-HD. Strikingly, we indeed identified a significant increase of the protein’s hydrodynamic radius with increasing 1,6-HD concentrations (Fig. 5f). As a control, the *R*_h_ of EGFP alone was monitored at the same 1,6-HD concentrations where no increase in *R*_h_ was observed. This expansion in protein size indeed suggests that 1,6-HD acts as a solvation agent for the FUS polypeptide chain and, likely, in particular for the intrinsically disordered regions of FUS. Hereby, the intramolecular interactions between amino-acid residues causative for protein compaction are disrupted by 1,6-HD and, hence, become less favourable in the presence of 1,6-HD, causing the 1,6-HD co-solvated protein to expand in size. Interestingly, this solvation effect is not limited to just the protein. A similar expansion was also observed for the PEG copolymer (see Supplementary Fig. S11). This highlights that 1,6-HD functions as a broadly acting solvation agent for biomolecules by decreasing their phase separation propensity by sequestering interactions.

## Significance and conclusions

Here, we have formulated a broadly applicable approach for the quantification of collective interactions capturing both compositional and energetic details of biomolecular phase separation by measuring dilute phase concentrations of one component only (Fig. 1 and 2). In establishing this framework, we have discovered that FUS phase separation at low salt concentrations is driven by the preferential exclusion of ions from condensates to decrease charge screening (Fig. 3). At high salt concentrations, in the reentrant regime, an opposing trend is observed, where ions preferentially partition into condensates to enable non-ionic interactions (Fig. 4). Moreover, our findings demonstrate that 1,6-HD disrupts condensates by functioning as a solvation agent, as confirmed by an expansion of the FUS polypeptide chain and the copolymer PEG with increasing concentrations of 1,6-HD (Fig. 5).

An important finding of our investigations into the thermodynamic driving forces of phase separation under the influence of salts is that even simple molecules can display switch-like partitioning and stabilisation depending on the environmental conditions. Specifically, in the low salt regime, KCl is largely preventative of phase separation as it apprehends protein interactions by charge screening. Therefore, phase separation occurs upon decreasing the salt concentration and, hence, it becomes favourable to exclude the ions from the dense phase. Conversely in the high salt regime, the increase of LiCl and CsCl salt concentration acts as the trigger for phase separation by driving non-ionic interactions, hence, favouring ion enrichment in the condensed phase. More generally, this might indicate that the partitioning of molecular species is largely a consequence of their propensity to enhance or counteract collective interactions of biomolecules.

Furthermore, our works provide unique insights into the mechanism of condensate modulation by 1,6-HD. 1,6-HD is one of the most commonly used phase separation modulators in the study of condensate stability, dissolution effects, and numerous other aspects^33,36–39^. It is broadly agreed to function as a hydrophobic disruptor^33^. Here, we reveal that this effect is a consequence of 1,6-HD acting as a solvating agent for biomolecules. Specifically, 1,6-HD decreases inter- and intermolecular interactions by favouring solvent interactions, as corroborated by an expansion of the polypeptide chain of FUS upon increasing the 1,6-HD. Given the important role of intrinsically disordered proteins in phase separation, this mechanism of action might translate to other systems where 1,6-HD dissolution effects are observed. In addition, our mechanistic study of 1,6-HD suggests that affecting the biomolecule native state and chain expansion is a powerful avenue for modulating phase behaviour. Hence, screening for molecules capable of inducing expansion or compaction of intrinsically disordered proteins might be a simple but effective drug discovery strategy.

Our work also pinpoints how quantification of collective interactions at different ionic strengths can inform on condensate modulation strategies. Our data highlight that FUS phase separation at low ionic strengths is driven by ion release from the polypeptide chain of the monomeric protein upon condensate formation. Hence, in a condensate modulation context, functional groups that are capable of complexing charged sites on the protein, such as carboxylic acids or highly charged functionalities, may likely constitute a promising modulation strategy to dissolve FUS condensates. The ion release mechanism from FUS condensates also points towards a potential dynamic control mechanism of phase separation in cells. Intracellular chloride ion levels can vary from a few millimolar to above a hundred millimolar^41^ and intracellular chloride levels can further be altered by changes in the expression levels of various ion channels, in the context of pathology^42,43^. Given the profound impact of ion partitioning on FUS phase separation, we speculate that changes in intracellular chloride ion concentrations could serve to modulate phase separation of FUS and possibly other proteins in cells.

Crucially, our approach presented here allows for the study of species, such as salt ions or small molecule compounds, which would otherwise be inaccessible by conventional labelling or detection strategies. This makes the assay especially suited to applications in more complex systems as the characterisation of the underlying driving forces can be performed by measuring the response of an individual component only. Here, the theoretical reduction framework alleviates the necessity to directly measure more components. The dilute-phase-centred analysis strategy presents a number of important advantages such as drastically decreasing material requirements as it circumvents the necessity to generate large amounts of dense phase volume. Dilute phase readouts can be generated easily by a multitude of commonly available measurement assays besides the confocal detection-based assay employed here.

Taken together, the quantification of collective interactions with the presented approach promises to be a powerful tool for studying phase separation in a wide range of contexts. We envision that the approach will find broad applicability in gaining mechanistic insights to rationally design condensate modulators, dissecting the impact of potential novel drug candidates, studying the impact of biologically relevant molecular species on *in vivo* phase transitions, and optimising designer phase separation systems for functional applications.

## Acknowledgements

The research leading to these results has received funding from Global Research Technologies, Novo Nordisk A/S (H.A., T.P.J.K.), the European Research Council under the European Union’s Horizon 2020 Framework Programme through the Marie Sklodowska-Curie grant MicroSPARK (agreement no. 841466; G.K.), the Herchel Smith Funds (G.K.), the Alzheimer Association Zenith Award (P.S.G.-H.) and the Wolfson College Junior Research Fellowship (G.K.). T.J.W. thanks the Harding Distinguished Postgraduate Scholar Programme.

## Author contributions

H.A., D.Q., G.K., T.J.W., and T.P.J.K. designed and conceptualised the study. H.A. and E.d.C. performed experiments. G.K., T.J.W., T.S., T.M.F., S.Q., N.A.E. and P.S.G.-H. provided materials and methods. H.A. and D.Q. analysed the data. H.A., D.Q., G.K., T.J.W., J.N.A., M.K., R.V.P. and T.P.J.K. interpreted data. H.A. and G.K. wrote the original draft of the paper. All authors discussed the results, commented on the manuscript, and contributed to the final manuscript.

## Data availability

The raw data and analysis code underlying this study will be made available upon request.

## References

1. Gallie, D. R. Protein-protein interactions required during translation. Plant Molecular Biology 50, 949–970 (2002).

2. Guarracino, D. A., Bullock, B. N. & Arora, P. S. Mini review: protein-protein interactions in transcription: a fertile ground for helix mimetics. Biopolymers 95, 1–7 (2011).

3. Pawson, T. & Nash, P. Assembly of cell regulatory systems through protein interaction domains. Science 300, 445–452 (2003).

4. Pawson, T. & Nash, P. Protein-protein interactions define specificity in signal transduction. Genes Dev 14, 1027–1047 (2000).

5. Banani, S. F., Lee, H. O., Hyman, A. A. & Rosen, M. K. Biomolecular condensates: organizers of cellular biochemistry. Nature Reviews Molecular Cell Biology 18, 285–298 (2017).

6. Shin, Y. & Brangwynne, C. P. Liquid phase condensation in cell physiology and disease. Science 357, eaaf4382 (2017).

7. Hyman, A. A., Weber, C. A. & Jülicher, F. Liquid-liquid phase separation in biology. Annual review of cell and developmental biology 30, 39–58 (2014).

8. Brangwynne, C. P., Tompa, P. & Pappu, R. V. Polymer physics of intracellular phase transitions. Nature Physics 11, 899–904 (2015).

9. Alberti, S., Gladfelter, A. & Mittag, T. Considerations and Challenges in Studying Liquid-Liquid Phase Separation and Biomolecular Condensates. Cell 176, 419–434 (2019).

10. Kedersha, N. & Anderson, P. Mammalian Stress Granules and Processing Bodies. in Methods in Enzymology vol. 431 61–81 (Academic Press, 2007).

11. Wolozin, B. & Ivanov, P. Stress granules and neurodegeneration. Nature Reviews Neuroscience 20, 649–666 (2019).

12. Guillén-Boixet, J. et al. RNA-Induced Conformational Switching and Clustering of G3BP Drive Stress Granule Assembly by Condensation. Cell 181, 346–361.e17 (2020).

13. Patel, A. et al. A Liquid-to-Solid Phase Transition of the ALS Protein FUS Accelerated by Disease Mutation. Cell 162, 1066–1077 (2015).

14. Alberti, S. & Dormann, D. Liquid-Liquid Phase Separation in Disease. Annu Rev Genet 53, 171–194 (2019).

15. Arter, W. E. et al. Rapid Generation of Protein Condensate Phase Diagrams Using Combinatorial Droplet Microfluidics. bioRxiv 2020.06.04.132308 (2020) doi:10.1101/2020.06.04.132308.

16. McCarty, J., Delaney, K. T., Danielsen, S. P. O., Fredrickson, G. H. & Shea, J.-E. Complete Phase Diagram for Liquid–Liquid Phase Separation of Intrinsically Disordered Proteins. J. Phys. Chem. Lett. 10, 1644–1652 (2019).

17. Villois, A. et al. Droplet Microfluidics for the Label-Free Extraction of Complete Phase Diagrams and Kinetics of Liquid–Liquid Phase Separation in Finite Volumes. Small n/a, 2202606 (2022).

18. Qian, D. et al. Tie-Line Analysis Reveals Interactions Driving Heteromolecular Condensate Formation. Phys. Rev. X 12, 041038 (2022).

19. Daoyuan Qian, Hannes Ausserwoger, Tomas Sneideris, Rohit Pappu, & Tuomas P. J. Knowles. Dominance metric in multi-component binary phase equilibria. bioRxiv 2023.06.12.544666 (2023) doi:10.1101/2023.06.12.544666.

20. Murakami, T. et al. ALS/FTD Mutation-Induced Phase Transition of FUS Liquid Droplets and Reversible Hydrogels into Irreversible Hydrogels Impairs RNP Granule Function. Neuron 88, 678–690 (2015).

21. Maharana, S. et al. RNA buffers the phase separation behavior of prion-like RNA binding proteins. Science 360, 918–921 (2018).

22. Qamar, S. et al. FUS Phase Separation Is Modulated by a Molecular Chaperone and Methylation of Arginine Cation-π Interactions. Cell 173, 720–734.e15 (2018).

23. Dormann, D. et al. ALS-associated fused in sarcoma (FUS) mutations disrupt Transportin-mediated nuclear import. The EMBO Journal 29, 2841–2857 (2010).

24. Dormann, D. & Haass, C. Fused in sarcoma (FUS): An oncogene goes awry in neurodegeneration. Molecular and Cellular Neuroscience 56, 475–486 (2013).

25. Tong, X. et al. Liquid–liquid phase separation in tumor biology. Signal Transduction and Targeted Therapy 7, 221 (2022).

26. Boija, A., Klein, I. A. & Young, R. A. Biomolecular Condensates and Cancer. Cancer Cell 39, 174–192 (2021).

27. Welsh, T. J. et al. Surface Electrostatics Govern the Emulsion Stability of Biomolecular Condensates. Nano Lett 22, 612–621 (2022).

28. Li, L. et al. Phase Behavior and Salt Partitioning in Polyelectrolyte Complex Coacervates. Macromolecules 51, 2988–2995 (2018).

29. Friedowitz, S. et al. Looping-in complexation and ion partitioning in nonstoichiometric polyelectrolyte mixtures. Science Advances 7, eabg8654.

30. Kozlov, A. G. et al. How Glutamate Promotes Liquid-liquid Phase Separation and DNA Binding Cooperativity of E. coli SSB Protein. Journal of Molecular Biology 434, 167562 (2022).

31. Suhling, K. et al. Fluorescence lifetime imaging (FLIM): Basic concepts and some recent developments. Medical Photonics 27, 3–40 (2015).

32. Suhling, K. et al. Imaging the Environment of Green Fluorescent Protein. Biophysical Journal 83, 3589–3595 (2002).

33. Krainer, G. et al. Reentrant liquid condensate phase of proteins is stabilized by hydrophobic and non-ionic interactions. Nature Communications 12, 1085 (2021).

34. Cacace, M. G., Landau, E. M. & Ramsden, J. J. The Hofmeister series: salt and solvent effects on interfacial phenomena. Quarterly Reviews of Biophysics 30, 241–277 (1997).

35. Kroschwald, S., Maharana, S. & Alberti, S. Hexanediol: a chemical probe to investigate the material properties of membrane-less compartments. Matters (2017) doi:10.19185/matters.201702000010.

36. Ambadipudi, S., Reddy, J. G., Biernat, J., Mandelkow, E. & Zweckstetter, M. Residue-specific identification of phase separation hot spots of Alzheimer’s-related protein tau. Chem. Sci. 10, 6503–6507 (2019).

37. Yamazaki, T. et al. Functional Domains of NEAT1 Architectural lncRNA Induce Paraspeckle Assembly through Phase Separation. Molecular Cell 70, 1038–1053.e7 (2018).

38. Strom, A. R. et al. Phase separation drives heterochromatin domain formation. Nature 547, 241–245 (2017).

39. Lu, H. et al. Phase-separation mechanism for C-terminal hyperphosphorylation of RNA polymerase II. Nature 558, 318–323 (2018).

40. Arosio, P. et al. Microfluidic Diffusion Analysis of the Sizes and Interactions of Proteins under Native Solution Conditions. ACS Nano 10, 333–341 (2016).

41. Kim, S., Ma, L., Unruh, J., McKinney, S. & Yu, C. R. Intracellular chloride concentration of the mouse vomeronasal neuron. BMC Neuroscience 16, 90 (2015).

42. Gururaja Rao, S., Patel, N. J. & Singh, H. Intracellular chloride channels: novel biomarkers in diseases. Frontiers in physiology 11, 96 (2020).

43. Peretti, M. et al. Chloride channels in cancer: Focus on chloride intracellular channel 1 and 4 (CLIC1 AND CLIC4) proteins in tumor development and as novel therapeutic targets. Biochimica et Biophysica Acta (BBA) -Biomembranes 1848, 2523–2531 (2015).

