## Supplementary Information for "Quantifying collective interactions in biomolecular phase separation"

* Contributed equally

### Materials and methods

#### Fabrication of microfluidic devices

Microfluidic devices were designed using AutoCAD software (Autodesk) followed by printing on acetate transparency masks (Micro Lithography Services). The replica master was obtained via standard soft-lithography steps using spin-coating of SU-8 photoresists (MicroChem) onto polished silicon wafers.[1] Typically, SU-8 3050 was applied to achieve a device height of approximately 100 µm. After UV exposure, using a custom-built LED-based apparatus[2], the precise heights of the features were measured using a profilometer (Dektak, Bruker). Devices were then produced in polydimethylsiloxane (PDMS). PDMS (Dow Corning) was mixed in a 10:1 (w/w) ratio with curing agent (Sylgard 184, Dow Corning) and poured onto the master, followed by degassing and baking for 1.5 h at 65°C. The PDMS was then removed from the master and punched using biopsy punches to generate inlet holes, after which the slab was bonded onto thin glass slides using oxygen plasma surface activation (Diener electronic, 40 % power for 30 s).

#### Reagents

Potassium chloride (KCl), Lithium Chloride (LiCl), Caesium Chloride, polyethylene glycol (PEG, 10 kDa), Tween 20, Tris(hydroxymethyl)aminomethane (TRIS), and 1,6-hexanediol (1,6-HD) were purchased from Sigma Aldrich (United Kingdom).

#### FUS protein expression

EGFP-tagged fused in sarcoma (FUS) was expressed as reported previously in an insect cell expression system[3]. After purification, the protein was stored in 50 mM TRIS buffer (pH 7.4), 500 mM KCl and at a total protein concentration of 70 µM.

#### EGFP synthesis and purification

pET28a vector harbouring the gene encoding His-tagged enhanced green fluorescent protein (EGFP) was purchased from GenScript. 40 mL of lysogeny broth (LB) medium supplemented with 36 µg/mL of kanamycin was inoculated with one colony of *E. coli* BL21 DE3 cells (New England Biolabs) harbouring the vector encoding EGFP. The cells were grown overnight at 37 °C and 180 rpm. The next day, 40 mL of fresh LB-kanamycin media was inoculated with 1 mL of overnight culture and grown for 2-3 hours, until the optical density reached ~0.5 optical units. Then, the cell culture was supplemented with 1 mM of isopropyl-beta-D-thiogalactopyranoside (IPTG, Sigma-Aldrich) and grown for additional 3 hours. The cells were harvested by centrifugation at 5000 × *g* and 4 °C for 10 min. Harvested cells were resuspended in lysis buffer (1 mg/mL lysozyme, 10 mM imidazole in PBS (pH 7.4)) and incubated on ice for 30 min. Subsequently, cells were sonicated on ice using 6 × 10 s on/pause cycles (at 42% power cycle). Sonicated cells were centrifuged for 30 min at 10.000 × *g* and 4 °C. The supernatant was collected and loaded onto 1 mL of equilibrated Ni-NTA slurry in a BioRad gravity flow column. The column was placed on a rocker and incubated in a cold room at 4 °C for 60 min. Subsequently, the column was placed in a ring stand, after the beads settled, the flow-through was let to run, then the column was washed 4 × 1 mL of wash buffer solution (20 mM imidazole in PBS (pH 7.4)). Subsequently, EGFP was eluted using 2 mL of elution buffer (250 mM imidazole in PBS (pH 7.4)). Collected EGFP solution was buffer exchanged to 50 mM TRIS buffer (pH 7.4), 150 mM KCl using 10 kDa cut-off centrifugal filters (Millipore).

#### Sample preparation and phase separation conditions

Phase separation was induced by gently mixing the individual components in Eppendorf tubes. Briefly, in a first step, buffer was added to the Eppendorf tube. The buffer was 50 mM TRIS (pH 7.4) supplemented with 0.01 % (v/v) Tween 20 in all cases. For FUS/KCl experiments, KCl stock solution (150 mM KCl in 50 mM TRIS (pH 7.4)) was then added to achieve the desired total KCl concentration, also considering the amount of KCl introduced from the protein stock. This was followed by the addition of FUS from the protein stock

(70 µM FUS-GFP in 50 mM TRIS buffer (pH 7.4), 500 mM KCl). For the FUS/KGlu experiments, the KCl concentration was kept constant at 40 mM, also considering the amount from protein addition later on. Then, KGlu stock solution (1000 mM KGlu in 50 mM TRIS, pH 7.4) was added to achieve the desired KGlu concentration prior to addition of FUS stock solution. For FUS/1,6-hexanediol and GUG aptamer experiments, 150 mM KCl and 15% (w/v) PEG (in 50 mM TRIS (pH 7.4), 150 mM KCl) were added to the buffer. Then either 1,6-hexanediol (50% (w/v) in 50 mM TRIS (pH 7.4), 150 mM KCl) or GUG aptamer (100 µM in 50 mM TRIS (pH 7.4), 150 mM KCl) were added, followed by addition of pre-diluted FUS stock (5 µM FUS-GFP in 50 mM TRIS buffer (pH 7.4), 150 mM KCl). This yielded solutions containing 5% (w/v) PEG with 7% (w/v) 1,6-hexanediol in 50 mM TRIS buffer (pH 7.4), 150 mM KCl in case of the 1,6-hexanediol experiment series or varying GUG concentrations between 1 and 1000 nM in GUG aptamer experiments. In all cases, typically 10 µL of sample was prepared and final FUS concentrations were 1 or 2 µM.

**Dilute phase concentration measurements using confocal detection**

Samples were prepared as described above and incubated for 5 minutes prior to the experiment. Microfluidic chips were filled with sample buffer followed by flushing the sample for 5 minutes at 100 µL/h to ensure equilibration of the channel (see Supplementary Fig. S12). To operate flow control through the microfluidic channel, negative pressure created by a glass syringe (Hamilton, Switzerland) and syringe pump (neMESYS, Cetoni, Germany) was applied. Thereby, channel loading was controlled by an inlet reservoir containing buffer or sample that was connected to the channel inlet. Following equilibration, dilute phase concentrations were measured using a home-built confocal setup. Differences between devices utilized were minimal compared to the baseline noise level (see Supplementary Fig. S13). The setup is equipped with picosecond-pulsed 485-nm and 640-nm laser sources, a 60x water-immersion objective (CFI Plan Apochromat WI 60x, NA 1.2, Nikon), and an automized sample stage onto which the microfluidic device can be mounted. An illustration of the setup can be found in Supplementary Fig. S2. Following laser excitation of the sample through the objective in a diffraction limited spot and collection of fluorescence through the same objective, emitted photons were registered via avalanche photo diodes and recorded using a time-correlated single-photon counting unit (HydraHarp400, Picoquant). Laser synchronization was done via a laser controller (Sepia PDL 828, Picoquant). The setup is further equipped with a lens-pinhole-lens system to remove out-of-focus light and with appropriate dichroic and bandpass filter sets to spectrally separate photons into blue- and red-emitting channels. The two lasers were run in pulsed-interleaved excitation (PIE) mode, driven by a synchronisation signal from the laser driver at a frequency of 25 MHz. Recording of data was done in T3 mode. Recorded macroscopic times $T$ were binned into 1-ms time intervals to give an intensity readout in units of number of photons per second. Signal collection from samples was typically performed for 30 s with collection times as low as 0.1 s being sufficient to calculate tie line gradients (see Supplementary Fig. S14). The intensity time traces generated in this manner typically display a stable baseline (see Fig. 2b, main text), which corresponds to the dilute phase concentrations, as the volume fraction of the dense phase can be assumed to be significantly smaller than the dilute phase. The baseline value can then be extracted from the maximum in the intensity histogram (see Fig. 2c, main text). The standard deviation of the baseline signal can be determined from numerical fitting to the lower intensity branch of the histogram. The higher intensity branch will show condensed species, as condensates exhibit high protein concentrations. Concentration conversion of the baseline intensities was performed by calibration via quadratic fitting to the intensity recorded of samples at different, known FUS concentrations under non-phase separated conditions.

**Fluorescence lifetime measurements**

Fluorescence lifetime decays were calculated by generating decay time histogram from a large number of photons. This was achieved by calculating the time difference between the APD signal and the last synchronisation event for each recorded photon. To estimate the lifetime of GFP in the dense phase, photons that correspond to the GFP fluorophore in the condensates are selected by applying a high-pass filter to the intensity readout previously computed by binning the macroscopic time $T$ in 1 ms intervals. More specifically, photons with a macroscopic time $T$ that falls in the high-intensity time window are identified as signal from condensates, and the $t$ values of all these photons are binned to give a single florescent decay histogram. To extract the lifetime, we truncated the histogram away from the instrument response function (IRF) and fitted an exponential decay function, whereas the inverse of the decay constant corresponds to the lifetime.

#### Microfluidic diffusional sizing

Microfluidic diffusional sizing was performed as described elsewhere[4]. Briefly, the Fluidity One-M instrument (Fluidic Analytics) was utilized for measurements. Auxiliary channels were first primed with 50 mM TRIS buffer (pH 7.4), 150 mM KCl. Subsequently, 3.5 µL of sample with 1 µM protein concentration and specified concentrations of 1,6-hexanediol were added. Each sample was measured three times. Detection was set for Alexa488 fluorescence detection and size-range settings of $R_H$ = 2–9 nm were applied on the Fluidity One-M instrument.

#### Determination of dilute phase response gradient

The dilute phase response gradient $R$ is determined directly from dilute phase line scan data. Line scans correspond to the change in dilute phase concentration of component A ($c_{\text{dil}}^{\text{A}}$) at constant total concentration of component A ($c_{\text{tot}}^{\text{A}}$ = const.) but varying total concentration of component B (vary $c_{\text{tot}}^{\text{B}}$). Here, the concentration of component B is varied from outside the two-phase coexistence region to well within the phase separated regime. This is repeated at two separate total concentrations of component A ($c_{\text{tot},1}^{\text{A}}, c_{\text{tot},2}^{\text{A}}$) to give two line scans. Both line scans are then individually fitted in the experimentally probed concentration range to give a function $c_{dil}^{A} = f(c_{\text{tot}}^{\text{B}})$. In the case of FUS and KCl (A = FUS, B = KCl), for example, the data shows a linear response to changes in KCl, thereby, the following form was chosen for phenomenological fitting to both line scans at $c_{\text{tot},1}^{\text{A}}$ and $c_{\text{tot},2}^{\text{A}}$.

$$c_{\text{dil}}^{\text{A}} = \begin{cases} c_{\text{tot}}^{\text{A}}, & c_{\text{tot}}^{\text{B}} > c_{\text{sat}}^{\text{B}} \\ \text{a} + \text{b} * c_{\text{tot}}^{\text{B}}, & c_{\text{tot}}^{\text{B}} < c_{\text{sat}}^{\text{B}} \end{cases}$$

In doing so, a, b and $c_{sat}^{B}$ can be determined for both line scans. The response gradient $R$ is then given as the average slope of both line scan fits at the phase boundary (i.e., $R = (\text{b}_1 + \text{b}_2)/2$). The dilute phase boundary gradient $P$ is then determined as follows:

$$P = \frac{c_{\text{tot},2}^{\text{A}} - c_{\text{tot},1}^{\text{A}}}{c_{\text{sat},2}^{\text{B}} - c_{\text{sat},1}^{\text{B}}} = \frac{\Delta c_{\text{tot}}^{\text{A}}}{\Delta c_{\text{sat}}^{\text{B}}},$$

With R and P in hand, the reduced tie line gradient $K$ as well as the component A dominance $D^{\text{A}}$ [5] is calculated:

$$\text{D}^{\text{A}} = \frac{\Delta f^{A}}{\Delta f} = \frac{K}{K-P} \text{ and } \text{K} = \frac{R*P}{R-P}.$$

For the sake of simplicity, the notation from the original theoretical description[5] were adjusted slightly, which are summarised in the Table below.

| **Notation here** | $P$ | $R$ | $D = \frac{R}{P}$ | $K = -P * \frac{D}{1-D}$ |
|---|---|---|---|---|
| **Reference notation**[5] | $-\frac{n_2}{n_1}$ | $R_2^1$ | $D^1 = -R_2^1 * \frac{n_1}{n_2}$ | $K_2^1 = \frac{n_2}{n_1} * \frac{D^1}{1-D^1}$ |

#### Phase diagram extrapolation

The phase boundary represents the set of conditions of dilute ($c_{\text{dil}}^{\text{A}}$, $c_{\text{dil}}^{\text{B}}$) and dense ($c_{\text{den}}^{\text{A}}$, $c_{\text{den}}^{\text{B}}$) phase concentrations formed as a result of the demixing of a set of total concentration conditions ($c_{\text{tot}}^{\text{A}}$, $c_{\text{tot}}^{\text{B}}$) within the two phase coexistence region. Here, the dilute phase part of the phase boundary can be extrapolated from component A dilute phase response function data only, as reported previously[6]. Briefly, for a specific condition ($c_{\text{tot},1}^{\text{A}}$, $c_{\text{tot},1}^{\text{B}}$) the experimentally determined dilute phase line-scans provide the corresponding dilute phase concentration $c_{\mathrm{dil,1}}^{\mathrm{A}}$. The dilute phase concentration of component B $c_{\mathrm{dil,1}}^{\mathrm{B}}$ can then be determined by following the tie line (with reduced tie line gradient K) originating in ($c_{\mathrm{tot,1}}^{\mathrm{A}}$, $c_{\mathrm{tot,1}}^{\mathrm{B}}$) until $c_{\mathrm{dil,1}}^{\mathrm{A}}$ (see Supplementary Fig. S4). Thereby, the point on the dilute phase branch of the phase boundary corresponding to the demixing of ($c_{\mathrm{tot,1}}^{\mathrm{A}}$, $c_{\mathrm{tot,1}}^{\mathrm{B}}$) can be simply written as:

$$(x,y) = (c_{\mathrm{dil,1}}^{\mathrm{A}}, c_{\mathrm{dil,1}}^{\mathrm{B}}) = (c_{\mathrm{dil,1}}^{\mathrm{A}}, c_{\mathrm{tot,1}}^{\mathrm{B}} - (c_{\mathrm{tot,1}}^{\mathrm{A}} - c_{\mathrm{dil,1}}^{\mathrm{A}})/\mathrm{K})$$

By assuming a constant tie line gradient and employing previously mentioned phenomenological fitting for the line-scans this can be repeated for any ($c_{\mathrm{tot,n}}^{\mathrm{A}}$, $c_{\mathrm{tot,n}}^{\mathrm{B}}$) within the experimentally characterised concentration range to determine the corresponding phase boundary condition ($c_{\mathrm{dil,n}}^{\mathrm{A}}$, $c_{\mathrm{dil,n}}^{\mathrm{B}}$).

#### 2D tie line reduction

A reduced tie line is by definition a collection of points on the 2-D experimental phase diagram that have the same dilute phase component A concentration (see Supplementary Fig. S1). When the full phase space is 2-D, the phase boundary and tie lines all lie on a plane, and near the dilute phase boundary points on each tie line all have the same dilute phase A concentration. As such, in the 2-D scenario the reduced tie line constructed using component A dilute phase concentration is equivalent to the higher dimensional tie line, and is also equivalent to the reduced tie line constructed using dilute phase B concentration. On the other hand, when the full phase space is 3-D, the phase boundary itself becomes a 2-D surface. The collection of points with the same component A dilute phase concentration constitutes another surface, since 3 (dimensions) - 1 (constraint) = 2 (degrees of freedom). This surface is a flat plane perpendicular to the component A concentration axis in the mixed region, where the dilute phase concentration is equivalent to the total concentration. When propagating this surface into the phase separated region, one has to bear in mind that many tie lines in the 3-D space share the same dilute phase A concentration, and these tie lines can be identified by the intersection between the trivial flat plane at $c_{\mathrm{tot}}^{\mathrm{A}}$ = const. and the 2-D phase boundary. We identify the tie lines of interest as the ones with one end on this intersection curve, and stitching them together produces a 2-D surface that divides the phase separates region into two parts. This newly generated surface can be thought of as a tie plane with respect to component A, and it is in general curved. When line scans performed in a 2-D experimental cross section of the full 3-D space, this cross-section intersects the tie plane and this intersection is the reduced tie line measured here. Hence, the reduced tie line depends on both tie lines and the phase boundary itself when more than 2 components are present.

### Supplementary Figures

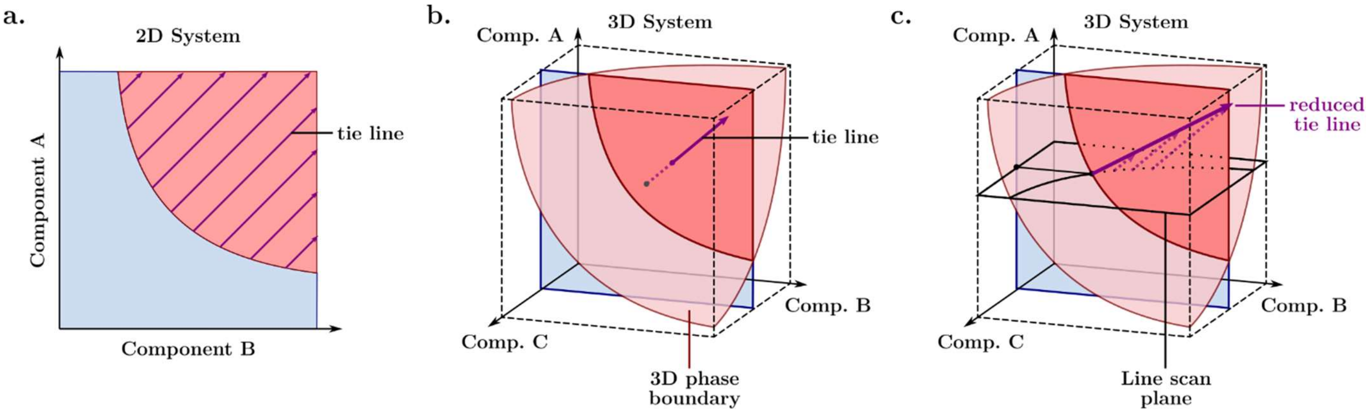


**Supplementary Figure S1. Schematic illustration of tie line reduction to the 2D measurement space. (a)** In a pure 2 component system, tie lines are constrained to the 2D measurement plane only. Thereby the reduced tie line gradient is equivalent to the 'higher' dimensional tie line. **(b)** Upon partitioning of additional components, not included in the measurement plane, tie lines become higher dimensional objects[5]. **(c)** Approximate information of the higher dimensional tie line gradient with respect to the measurement plane of interest can then be extracted from tie line reduction. The reduced tie line then represents the intersections of tie lines originating from a fixed component A dilute phase concentration with the measurement plane[5].

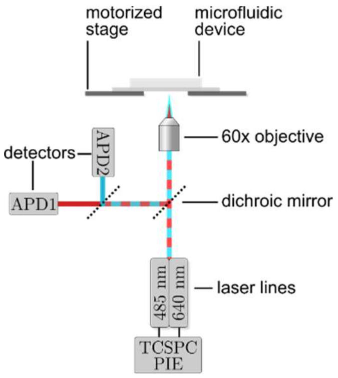


**Supplementary Figure S1. Schematic representation of the custom-built two-colour confocal measurement setup.** The setup consists of two picosecond-pulsed laser lines for 485-nm and 640-nm excitation, operated in pulsed interleaved excitation (PIE) mode. The laser light excites molecules in a microfluidic device through a high magnification objective. Emitted photons are spectrally separated and then detected by avalanche photodiodes (APDs) and time-correlated single-photon counting (TCSPC) electronics.

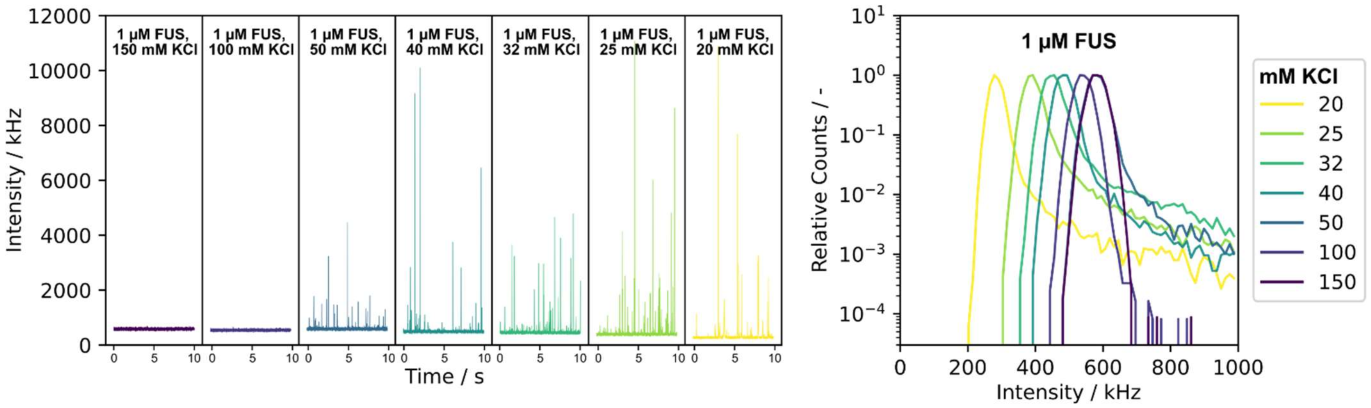


**Supplementary Figure S2. Time trace and intensity histogram line scan data for FUS/KCl. (left panels)** Representative individual intensity time traces for FUS at 1 uM (2 uM shown in main text Fig 2.) and varying KCl concentrations recorded using a microfluidic flow cell connected to confocal detection unit. Time traces show appearances of larger numbers of intensity bursts and decrease in the baseline intensity due to phase separation. Time traces measurements were performed in triplicates with additional repeats not shown (see SI Fig. 12 for exemplary variation between devices). **(right panel)** Intensity histograms of recorded time traces, with the maximum of the distribution representing the dilute phase concentration. These individual maxima are plotted against the total KCl concentration to give dilute phase line scan data.

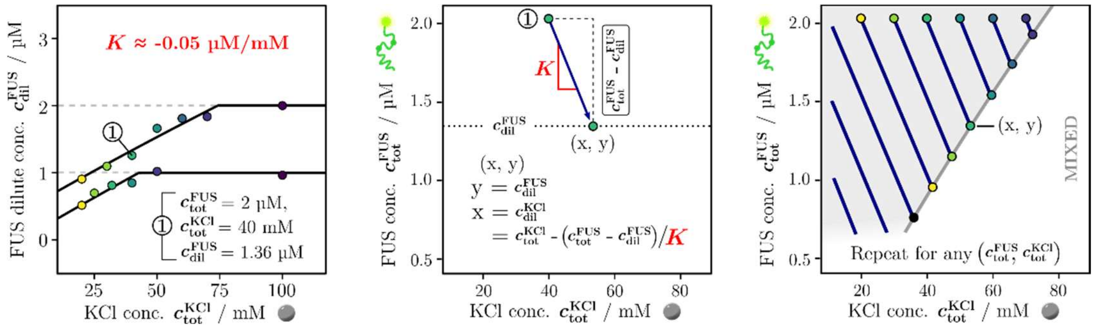


**Supplementary Figure S3. Phase boundary extrapolation from dilute phase concentration response functions on the example of FUS/KCl. (a)** Points on the phase boundary are identified by using the experimentally determined dilute phase concentration response functions and tie line gradient (left panel). Dilute phase response functions and determination of tie line gradients alone are sufficient to extrapolate the phase diagram in the measured concentration range. **(b)** A point on the phase boundary is then identified by intersecting tie lines and dilute phase concentration of a given total concentration point (1). Points on the phase diagram are identified by intersecting the measured dilute phase concentration and the origin of the tie line gradient.

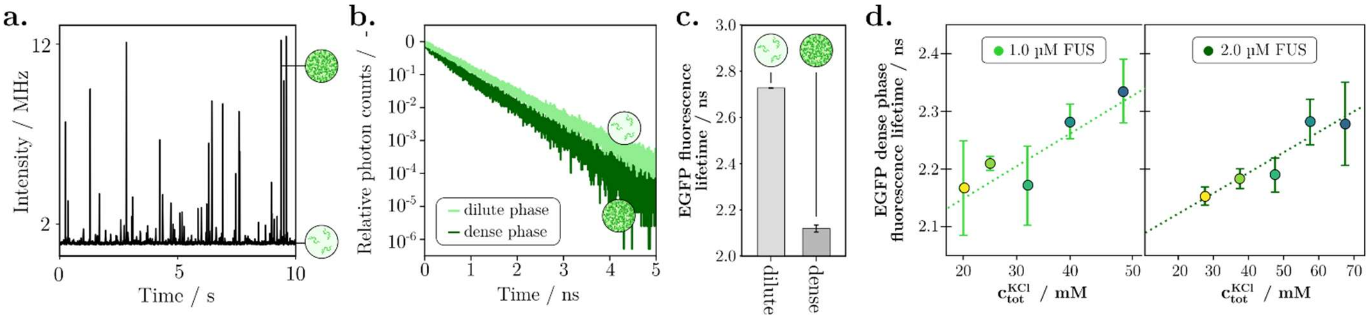


**Supplementary Figure S4. Dense phase fluorescent lifetime determination.** **(a)** Intensity time trace of 20 mM KCl and 1 µM FUS sample injected into microfluidic channel connected to confocal detection unit. Intensity bursts stem from the protein condensed phase, whereas the dilute phase forms a stable baseline. **(b)** Histogram of photon arrival times post excitation for dense and dilute phase signals, with the dense phase showing a faster decay. **(c)** Comparison of calculated dense and dilute phase fluorescent lifetimes from **(b)**. **(d)** Dense phase fluorescence lifetimes with changing KCl concentrations at both 1 and 2 µM FUS.

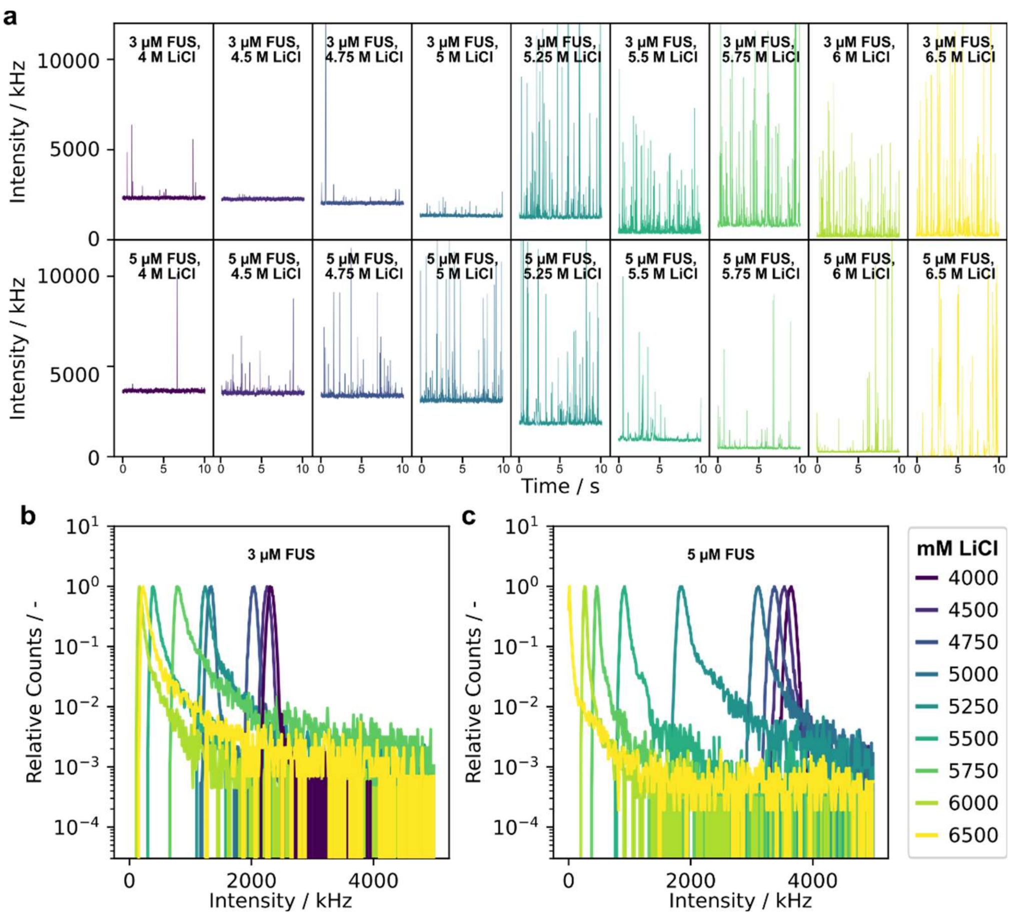


**Supplementary Figure S5. Time trace and intensity histogram line scan data for FUS/LiCl in the high salt reentrant regime.** **(a)** Representative individual intensity time traces for FUS at 3 and 5 uM and varying LiCl concentrations. Time traces measurements were performed in triplicates with additional repeats not shown. **(b, c)** Intensity histograms of recorded time traces, which are used to construct dilute phase line scan data at 3 and 5 uM FUS.

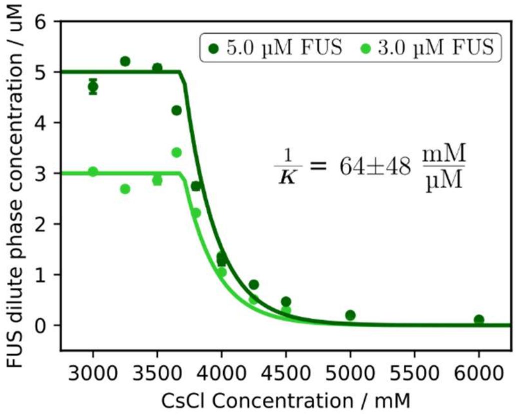


**Supplementary Figure S6. Tie line gradient analysis of FUS and CsCl. (a)** FUS dilute phase concentration change against total CsCl concentration and determination of the tie line gradient.

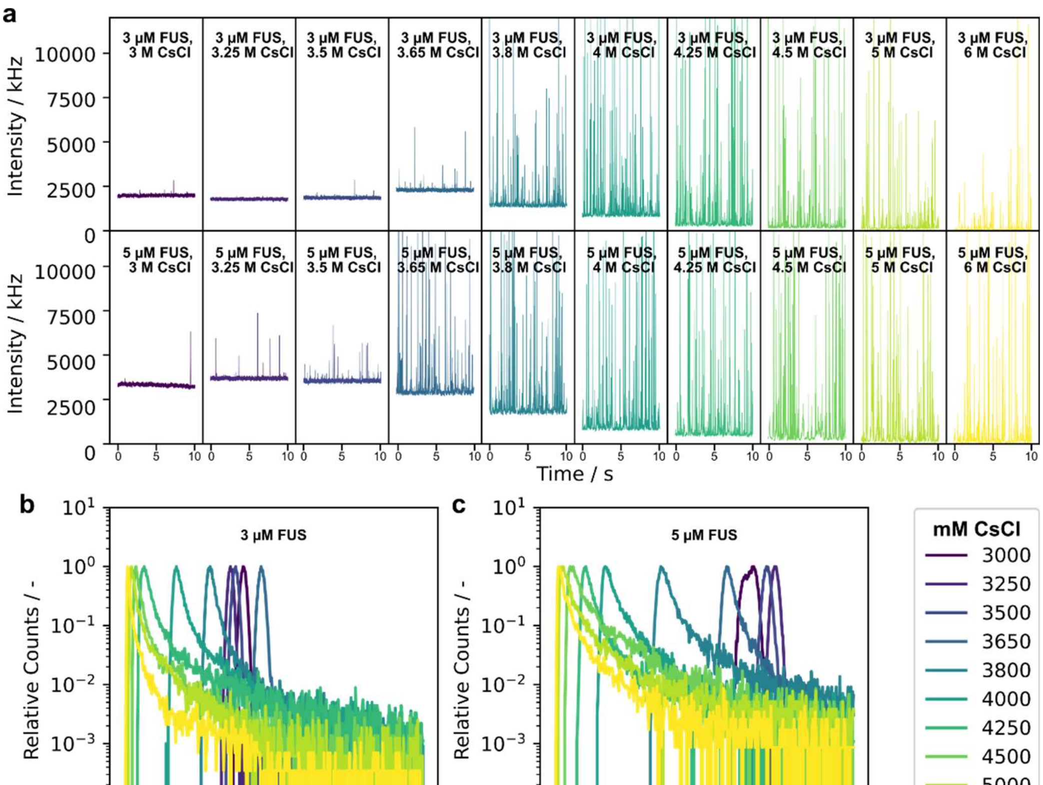


**Supplementary Figure S7. Time trace and intensity histogram line scan data for FUS/CsCl in the high salt reentrant regime. (a)** Representative individual intensity time traces for FUS at 3 and 5 uM and varying CsCl concentrations. Time traces measurements were performed in triplicates with additional repeats not shown. **(b, c)** Intensity histograms of recorded time traces, which are used to construct dilute phase line scan data.

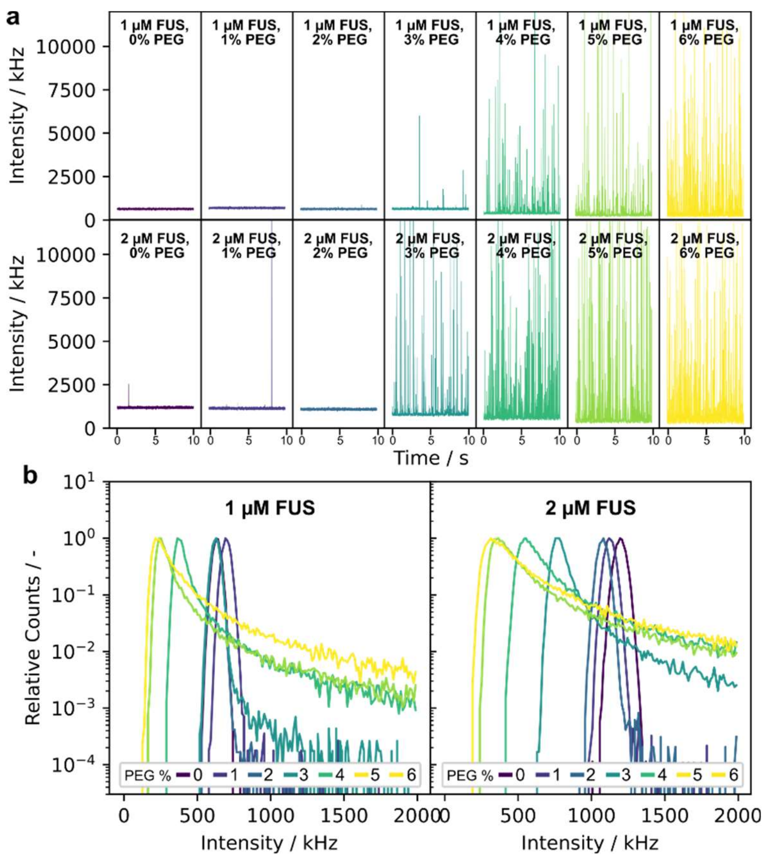


**Supplementary Figure S8. Time trace and intensity histogram line scan data for FUS/PEG. (a)** Representative individual intensity time traces for FUS at 1 and 2 uM and varying PEG concentrations. Time traces measurements were performed in triplicates with additional repeats not shown. **(b)** Intensity histograms of recorded time traces, which are used to construct dilute phase line scan data.

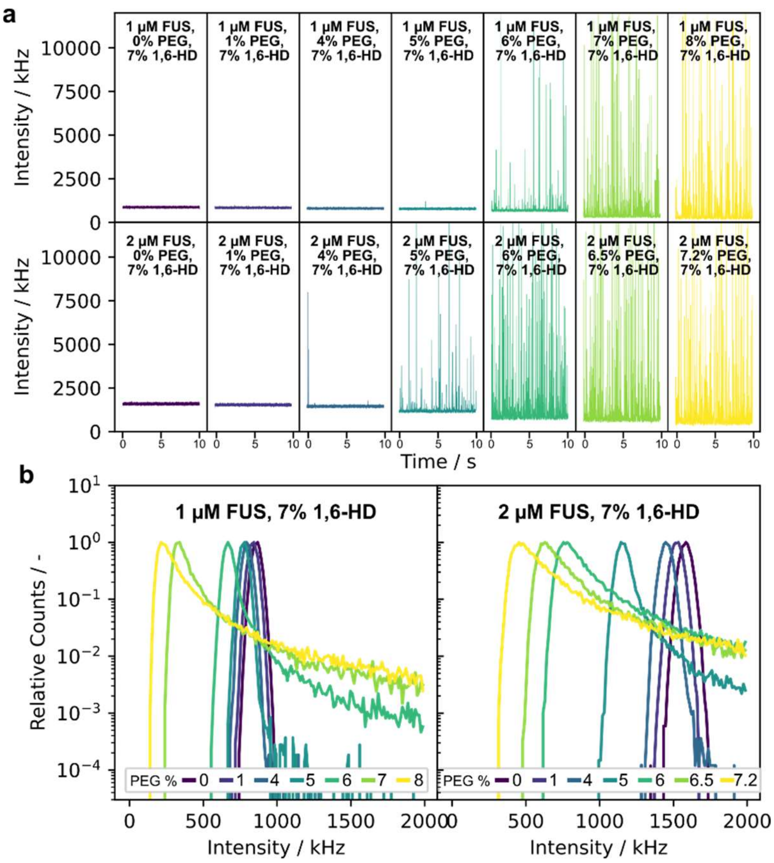


**Supplementary Figure S9. Time trace and intensity histogram line scan data for FUS/PEG in presence of 1,6-HD. (a)** Representative individual intensity time traces for FUS at 1 and 2 uM and varying PEG concentrations at 7 (w/v)% 1,6-hexanediol. Time traces measurements were performed in triplicates with additional repeats not shown. **(b)** Intensity histograms of recorded time traces, which are used to construct dilute phase line scan data.

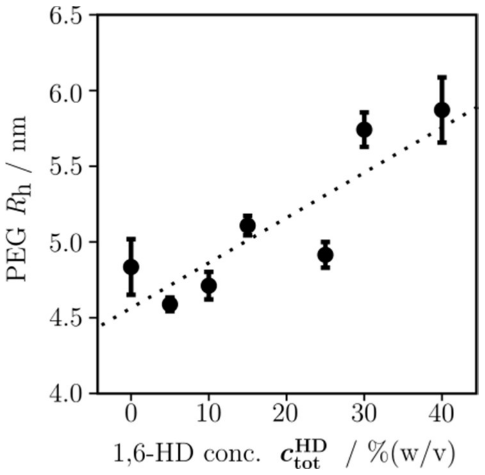


**Supplementary Figure S11. PEG expansion under influence of hexanediol.** Changes in hydrodynamic radius $R_h$ of fluorescently labelled PEG (20 kDa) at 0.1 (w/v)% as a function of increasing 1,6-HD concentration.

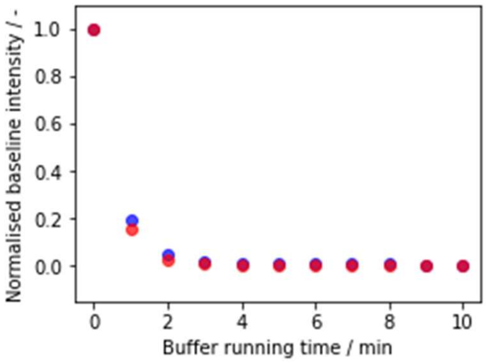


**Supplementary Figure S12. Analysis of necessary channel equilibration times after the change of sample.** Baseline intensity in the blue-emitting (blue) and red-emitting (red) channels. Shown is the normalized for the maximum intensity at the start of washing versus the runtime of the buffer washing. The microfluidic channel was first equilibrated with 1 µM FUS, 10 nM GUG aptamer and 8% (w/v) PEG at 50 mM TRIS (pH = 7.4), 150 mM KCl and subsequently flushed with 50 mM TRIS (pH = 7.4), 150 mM KCl at 100 µL/h to estimate the necessary channel washing/equilibration time. After 3 minutes of buffer wash, 99.3 % of the initial total intensity are removed.

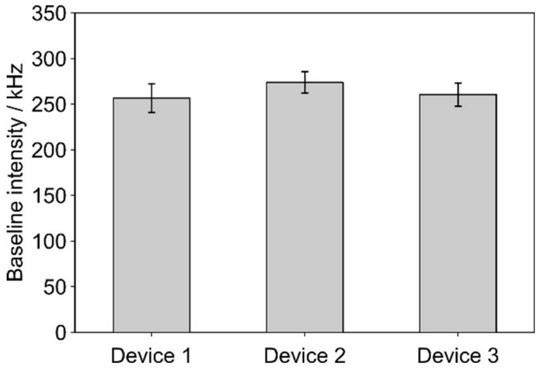


**Supplementary Figure S13. Variations in dilute phase concentration measurements between devices.** Baseline intensity of FUS at 1 µM and 5% (w/v) PEG in 3 different devices. Variations between devices are within measurement error.

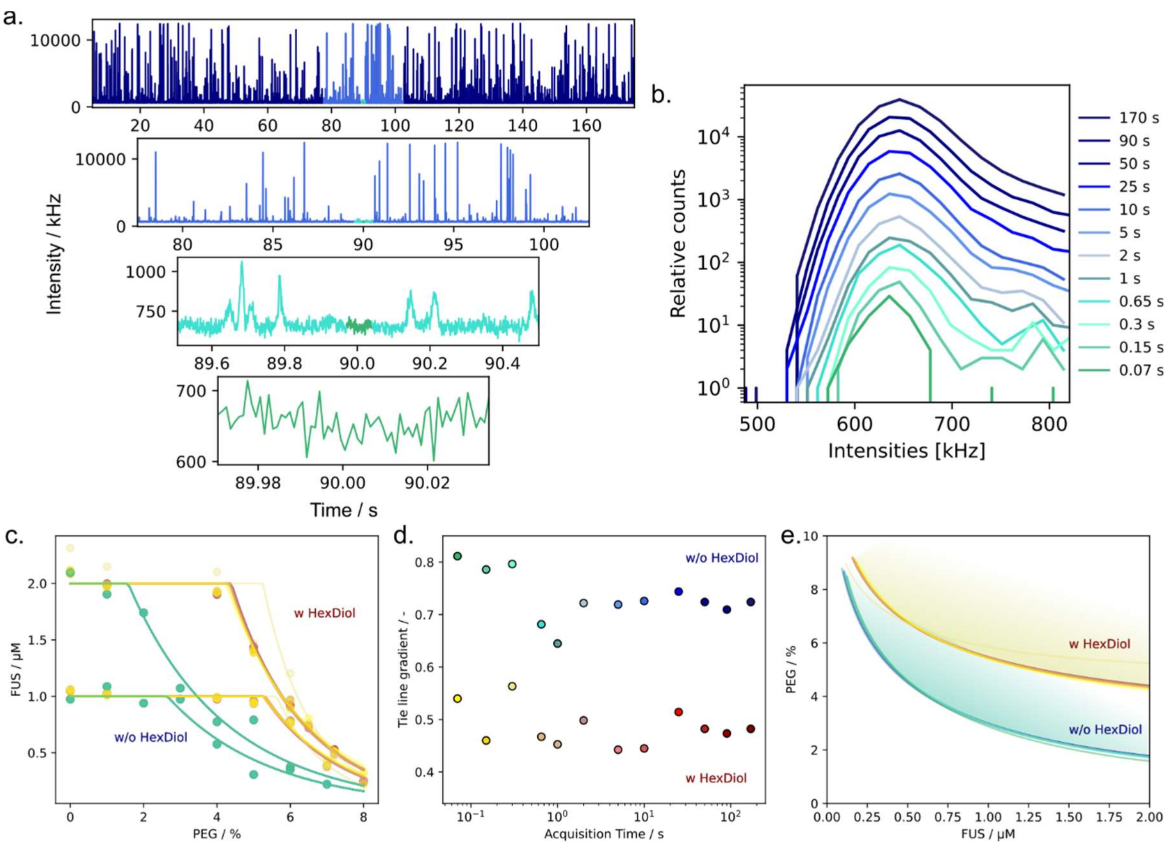


**Supplementary Figure S14. Analysis of the necessary acquisition time for obtaining accurate readouts of dilute phase concentrations.** **(a)** Zoom in of exemplary time traces acquired with confocal detection from total time spanning 180 down to 0.05 s. **(b)** Histograms of all intensity entries recorded in the measurement for decreasing total acquisitions times. **(c)** Overlay of FUS dilute phase concentrations obtained from different acquisition time windows for increasing PEG percentages. **(d)** Re-analysis of the tie line gradient of the FUS/PEG systems with and without 1,6-hexanediol using different total acquisition times. Color code: dark blue to turquoise and dark red to yellow depict the acquisition time axis **(e)** Overlay of phase boundaries obtained from different acquisition time windows.

### Supplementary References

1. Duffy, D. C., McDonald, J. C., Schueller, O. J. A. & Whitesides, G. M. Rapid Prototyping of Microfluidic Systems in Poly(dimethylsiloxane). *Anal. Chem.* **70**, 4974–4984 (1998).
2. Challa, P. K., Kartanas, T., Charmet, J. & Knowles, T. P. J. Microfluidic devices fabricated using fast wafer-scale LED-lithography patterning. *Biomicrofluidics* **11**, 014113 (2017).
3. Patel, A. *et al.* A Liquid-to-Solid Phase Transition of the ALS Protein FUS Accelerated by Disease Mutation. *Cell* **162**, 1066–1077 (2015).
4. Arosio, P. *et al.* Microfluidic Diffusion Analysis of the Sizes and Interactions of Proteins under Native Solution Conditions. *ACS Nano* **10**, 333–341 (2016).
5. Daoyuan Qian, Hannes Ausserwoger, Tomas Sneideris, Rohit Pappu, & Tuomas P. J. Knowles. Dominance metric in multi-component binary phase equilibria. *bioRxiv* 2023.06.12.544666 (2023) doi:10.1101/2023.06.12.544666.
6. Qian, D. *et al.* Tie-Line Analysis Reveals Interactions Driving Heteromolecular Condensate Formation. *Phys. Rev. X* **12**, 041038 (2022).
